# Detecting (un)seen change: The neural underpinnings of (un)conscious prediction errors

**DOI:** 10.1101/832386

**Authors:** Elise G. Rowe, Naotsugu Tsuchiya, Marta I. Garrido

## Abstract

Detecting changes in the environment is fundamental for our survival. According to predictive coding theory, detecting these irregularities relies both on incoming sensory information and our top-down prior expectations (or internal generative models) about the world. Prediction errors (PEs), detectable in event-related potentials (ERPs), occur when there is a mismatch between the sensory input and our internal model (i.e., a surprise event). Many changes occurring in our environment are irrelevant for survival and may remain unseen. Such changes, even if subtle, can nevertheless be detected by the brain without emerging into consciousness. What remains unclear is how these changes are processed in the brain at the network level. Here, we used a visual oddball paradigm, in which participants engaged in a central letter task during electroencephalographic (EEG) recordings while presented with task-irrelevant high- or low-coherence background, random-dot motion. Critically, once in a while, the direction of the dots changed. After the EEG session, we confirmed that changes in motion direction at high- and low-coherence were visible and invisible, respectively, using psychophysical measurements. ERP analyses revealed that changes in motion direction elicited PE regardless of the visibility, but with distinct spatiotemporal patterns. To understand these responses, we applied Dynamic Causal Modelling (DCM) to the EEG data. Bayesian Model Averaging showed visible PE relied on a release from adaptation (repetition suppression) within bilateral MT+, whereas invisible PE relied on adaptation at bilateral V1 (and left MT+). Furthermore, while feedforward upregulation was present for invisible PE, the visible change PE also included downregulation of feedback between right MT+ to V1. Our findings reveal a complex interplay of modulation in the generative network models underlying visible and invisible motion changes.

## INTRODUCTION

Detecting changes in the environment is fundamental for our survival because these may indicate potential rewards or threats (LeDoux, 1996; Panksepp, 1998; Rensink, 2002). According to predictive coding theory, we detect these irregularities by comparing incoming (bottom-up) sensory information with an internal model based either on prior sensory information or on (top-down) beliefs or expectations (Friston, 2005; Friston & Stephan, 2007; Hohwy, 2013; Clark, 2013). Mismatch between the sensory input and our internal model results in a surprise event, signalled by a prediction error (PE) response in event-related potentials (ERPs). These PE responses represent the difference in the neural activation between expected (frequent or ‘standard’) and unexpected (rare/surprising or ‘deviant’) events and have been used extensively in the auditory modality to yield the mismatch negativity (MMN; Näätänen et al., 1978; Näätänen 1992; Garrido et al., 2007). Over the last decade, protocols for inducing visual PE (or visual MMN, ‘vMMN’, more specifically) have been developed, with a focus on situations where individuals are aware of a change (Stefanics et al., 2014; Pazo-Alvzrez et al., 2003; Kremláček et al., 2016). In our everyday lives, however, many changes in the sensory environment that evoke PEs may go unnoticed (Czigler, 2014; Stefanics et al., 2014). What remains unclear is how such sensory changes are processed by the brain at the network level.

One of the advantages in investigating non-conscious PE using visual, compared to auditory, stimuli is a richer set of psychophysical methods available to render otherwise salient visual stimuli invisible to participants (Faivre et al., 2017). Employing such tools, researchers have demonstrated PE responses to changes without conscious awareness of these changes. These PE responses in EEG include those using backward masking (Czigler et al., 2007; Kogai et al., 2011), binocular rivalry (Jack et al., 2015; 2017), rapid unmasked presentation (1 ms, Bernat et al., 2001, and 10 ms; Brazdil et al., 2001), and attentional blink (Berti, 2011). The researchers isolated PE ERPs by comparisons between standard and deviant trials, finding stronger and more widespread activation associated with conscious than non-conscious visual PE. ERP analyses alone, however, cannot inform us of the underlying neurocircuitry underpinning the PE to changes that do, and do not, emerge into consciousness.

Dynamic Causal Modeling (DCM) is a technique useful for inferring the underlying causal network of dynamical systems (Friston, 2003). Boly et al., (2011), applied DCM to auditory PEs in EEG recorded from healthy participants and two populations of brain damaged patients: minimally conscious and vegetative state patients to show that feedback connectivity (or top-down modulation) was reduced in unconscious vegetative patients compared to minimally conscious patients (and healthy controls). This study suggests a potential involvement of feedback modulations in regulating the level of conscious awareness of PE generating stimuli. However, no studies have performed network level examination of non-conscious processing in fully awake healthy participants and whether this also abolishes top-down feedback connections.

To understand the differences in the neural circuitry between sensory changes that can and cannot be consciously perceived, we aimed to elicit visual PE using visible and invisible changes in motion direction. To achieve the desired level of visibility of motion stimuli, we manipulated the coherence of random-dot motion. We used DCM to estimate effective connectivity and examined the involvement of feedback connections for visible and invisible changes.

## MATERIALS AND METHODS

### Participants

Twenty-eight healthy university students participated in this study (10 females, aged 18-40, *M* = 22.44, *SD* = 4.30). All participants reported no history of neurological or psychiatric disorder or previous head trauma resulting in unconscious comatose states. All participants gave written informed consent in accordance with the guidelines of the University of Queensland’s ethics committee.

### Experimental design

Continuous EEG data were recorded during the main task using a Biosemi Active Two system with 64 Ag/AgCl scalp electrodes arranged according to the international 10-10 system. Data were recorded at a sampling rate of 1024 Hz. After the EEG was setup, participants practiced the 1-back letter task for 1 min (see below) before being tested in the main task. The main task was followed by psychophysical tests to assess the discriminability of motion stimuli (see below). We did not record EEG during these follow-up psychophysical tests. The entire experiment took under two hours, including the setup of the EEG.

### Display

The experiment was performed using a Macbook Pro and external screen with a resolution of 1920 x 1080 pixels with a 60 Hz refresh rate. All participants were seated at a viewing distance of 50 cm (without a chinrest). The experiments were programmed using the Psychophysics Toolbox extensions (Brainard, 1997; Pelli, 1997; Kleiner et al., 2007) for MATLAB (version 2014b).

### Task and visual stimuli during EEG measurement

To achieve the desired level of visibility and to induce visual PE to visible and invisible changes, we manipulated the coherence of random-dot motion (Britten et al., 1992). In short, it is well-known that the direction of motion can be consciously discriminated when the level of motion coherence of random dots is sufficiently high. As the level of coherence approaches zero, participants can no longer consciously see the direction of motion (Watanabe et al., 2011). We exploited this useful feature of motion perception and designed a paradigm to elicit prediction errors from visible and invisible changes in the motion direction of a cloud of random dots (Figure 1, demo: https://figshare.com/s/76484519f510ba74891b). These motion stimuli were never task-relevant during the main EEG experiment. Instead, participants were instructed to focus on a central 1-back task and be as quick and accurate as possible.

**Figure 1.**
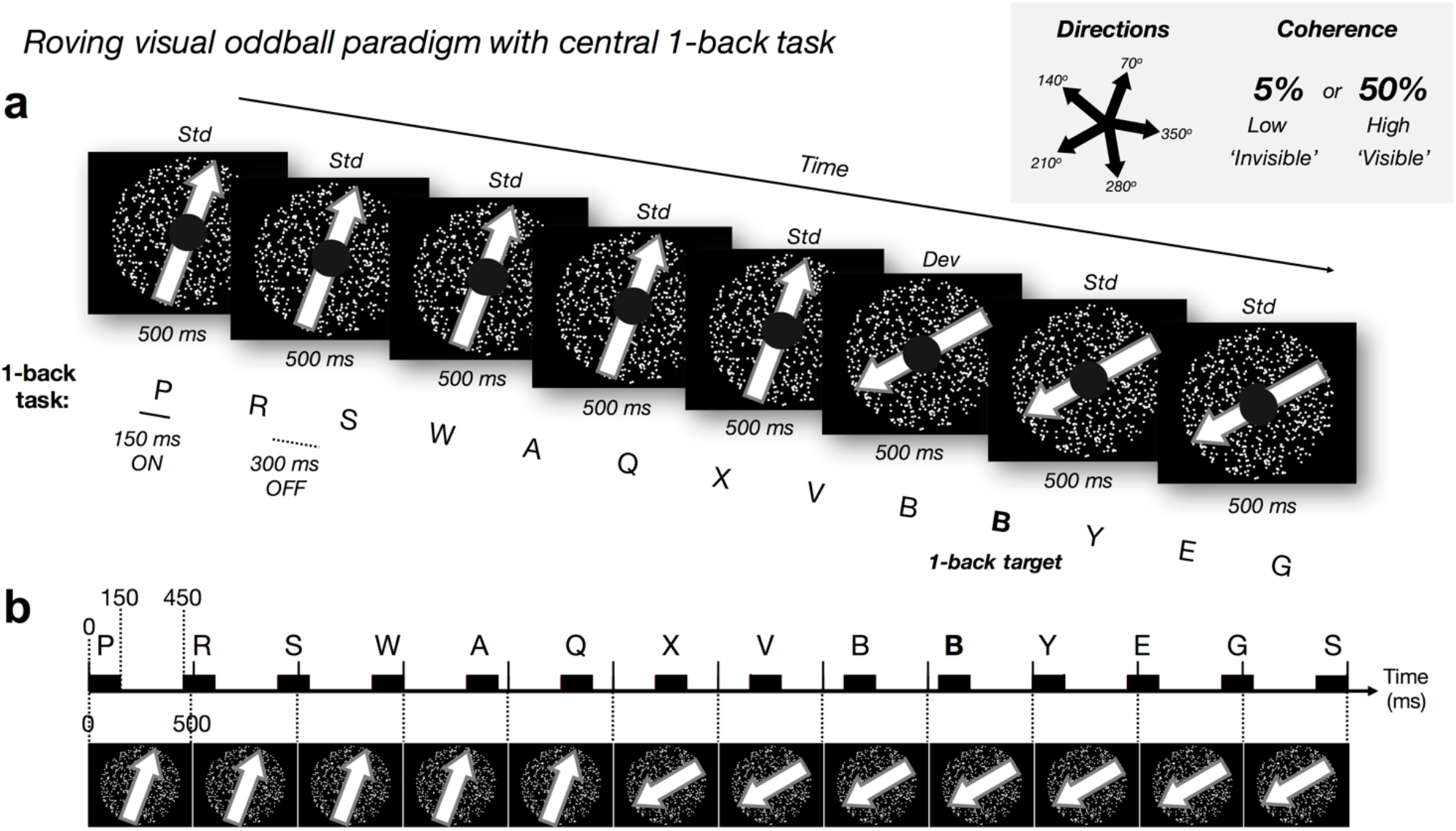
Experimental paradigm. (a) Roving visual oddball paradigm with dot motion at five possible directions and two possible coherence levels (low: 5% and high 50%). We manipulated *motion coherence*: high and low, and *motion direction*: standard (‘std’, frequent: no change) and deviant (‘dev’, rare/surprising: change of direction). The ‘roving design’ meant each deviant motion direction became the new standard direction over repetitions. We presented the task-irrelevant random-dot motion (500 ms per trial) in the visual periphery. Every 5 to 8 ‘standard’ trials the global direction of the dots changed (the ‘deviant’; randomly selected direction). Every 26 to 30 trials (i.e. 13-15 seconds), we changed both the direction and coherence level of the motion (but discarded EEG events and behavioural responses to these ‘double deviants’). Participants focused on a central 1-back letter task (150 ms ON and 300 ms OFF: desynchronized with most of the motion trials, aligning once every nine trials), responding when the same letter was repeated in succession (e.g., bold letter, ‘B’). Participants were instructed to ignore the motion stimuli and to be as fast and as accurate in the central task. (b) Schematic illustration of the timecourse for motion trials (every 500 ms) and 1-back letter presentation (black rectangles represent 150 ms letter ON).

Our motion stimuli comprised 750 white dots enclosed within a 30° diameter circular area on a black background. During each 500 ms trial, each dot moved in a straight trajectory at 14°/sec. When any dot went outside of the stimulus window, the dot was redrawn in the opposite location to keep the density of the dots constant. At the first frame of each 500 ms trial, we removed all dots and redrew them at new random locations at the same time (i.e. no blank interval between trials). Please refer to the exemplar video provided for further clarity of the stimuli.

To elicit PE responses, we utilised a roving visual oddball paradigm (i.e. 5-8 repetitions before a change) with dot motion at 5 possible non-cardinal directions (70° 140°, 210°, 280° and 350°; with 0° representing motion to the right) and 2 possible coherence levels (high: 50% and low 5%). Motion direction was manipulated in a 2 x 2 design comparing the *coherence* levels: high and low, and *motion direction*: standard (frequent: no change) and deviant (rare/surprising: change of direction). Mean number of low-coherence standard and deviant trials was 829 (SD = 65) and 136 (SD = 12), respectively. Mean number of high-coherence standard and deviant trials was 832 (SD = 64) and 136 (SD = 13), respectively.

One experimental block comprised 180 trials. At the beginning of the block, both the direction and the coherence of the motion was chosen randomly from the five possible directions and two coherence levels. The direction and coherence level remained unchanged for 5 to 8 trials before we changed the global motion direction randomly into one of the other 4 directions. Throughout the paper, we refer to the trials with unchanged motion direction as the standard trials and to the trials in which the direction changed as the deviant trials. Each deviant trial was then followed by another 5 to 8 trials without a change in motion direction, which we considered as the new standard trials. In some deviant trials, (once in every 26 to 30 trials), we also changed the coherence level. We did not include any of these ‘double deviant’ trials in our EEG or behavioural analysis.

At the centre of the visual display, we presented a 1-back task within a 5° diameter black circle. Here, a sequence of randomly selected white letters (~1.8° visual angle in size) was presented every 450 ms (i.e. 150 ms on and 300 ms off). The first of these letter stimuli was presented at the onset of the first motion trial but became (mostly) desynchronized in time after this (except for every 9 motion trials where the onset times realigned, see Figure 1b). Participants were required to attend to this letter stream and press the spacebar as quickly as possible whenever they detected the repetition of any letter (which could occur every 10 to 15 letters; Figure 1b). On average, the number of letter repeat targets per participant across the experiment was 191 (range from 177 to 196, SD = 3.69).

#### Behavioral analysis on the 1-back task during the EEG measurement

We examined participants’ accuracy during high- and low-coherence motion, defined as the hit and false alarm rates (and the sensitivity measure of d’; Green & Swets, 1966). For this, we defined a ‘hit’ as a response made between 200 to 1,000 ms after the onset of a letter repeat. Responses outside of this were considered a false alarm. Due to the fast presentation rate of our target events, we chose to use the method described by Bendixen & Andersen (2012; ‘Method E’, pp. 930) to calculate our false alarm rate. Here, the number of non-target events (the denominator for calculating the false alarm rate) were defined as the duration of the experiment divided by the average time it would take to execute a response (i.e. 300 ms) minus the number of target events. In using this method, we aimed to overcome the bias in using traditional signal detection theory methods in paradigms with high event rates. Two participants failed to correctly detect above 50% of the targets across the entire experiment. We removed these participants from further analyses.

#### Follow-up psychophysics tasks

To ensure the visibility and invisibility of high- and low-coherence motion direction changes, respectively, we tested each participant’s performance with two follow-up psychophysics tasks. The first task required participants to make a direction discrimination at high- or low-coherence levels. The second psychophysics task, performed in a subset of participants (N=8 out of 28 participants), required them to report when they ‘felt’ or ‘sensed’ a change in direction occurred at both high- or low-coherence levels.

#### Follow-up psychophysics task 1: direction discrimination task

To estimate the discriminability of the direction of motion, we ran a four alternative-forced choice (4AFC) direction discrimination task based on a study design with similar stimuli to ours (Tsushima et al., 2006). To conservatively estimate the discriminability, we reduced the direction alternatives to 4 possibilities (80°, 160°, 240°, and 320°), which were different from the motion directions used in the main EEG experiment in order to reduce the chance of perceptual learning which can occur even to sub-threshold dot motion (Watanabe et al., 2011; Tsushima et al., 2006). Furthermore, we aimed to avoid any habituation effects that may have arisen if we had used the same motion directions in both experiments. These effects, in turn, could have improved or reduced task performance. Rather than inducing these uncertain results, we opted for slightly different stimuli as an alternative solution. In each trial, we presented the motion for 915 ms without the central letter task (i.e. the small, black central circle remained but the letters were absent). We selected this extended presentation time based on the study by Tsushima et al., (2006) and in order to conservatively estimate the motion direction discriminability. We did not redraw all the dots after 500 ms as in the EEG experiment but, instead, the dots remained moving for 915 ms (except when it came to the boundary, see above). We randomly selected a motion direction in each trial and pseudo-randomly intermixed 4 coherence levels (2.5%, 5%, 25% and 50%) in equal proportions across 120 trials. We incorporated two additional motion coherence levels to reduce the chance of implicit learning based on the motion coherence level. At the end of every trial, participants reported the perceived direction of motion from the 4 possible alternatives.

### Behavioral results psychophysics task 1: *Confirming visibility and invisibility of high- and low-coherence motion direction changes in each individual*

We used performance in our follow-up psychophysics direction discrimination task to confirm that high- and low-coherence motion direction changes were visible and invisible, respectively. Based on the performance, we excluded participants from the following EEG analysis based on two criteria: 1) performing above chance at low (5%) coherence motion condition or 2) performing below chance at high (50%) coherence motion condition (Figure 2a and 2b). To estimate if the observed discrimination accuracy was above chance, we performed a bootstrap analysis (with replacement) over 1,000 repetitions per participant. For each participant, at a given coherence level, we generated a surrogate sequence of correct vs incorrect discriminations across 30 trials (e.g. correct, incorrect, correct, correct, correct….) which reflected the proportion of correct responses at the number of trials we tested. We then used 2.5%ile and 97.5%ile of the bootstrapped distribution as the confidence interval. That is, for the low (5%) coherence condition, if the bottom end of the confidence interval was above chance (25% correct), we assumed that the participant may have been aware of the direction of motion during the EEG experiment (Figure 2b).

**Figure 2.**
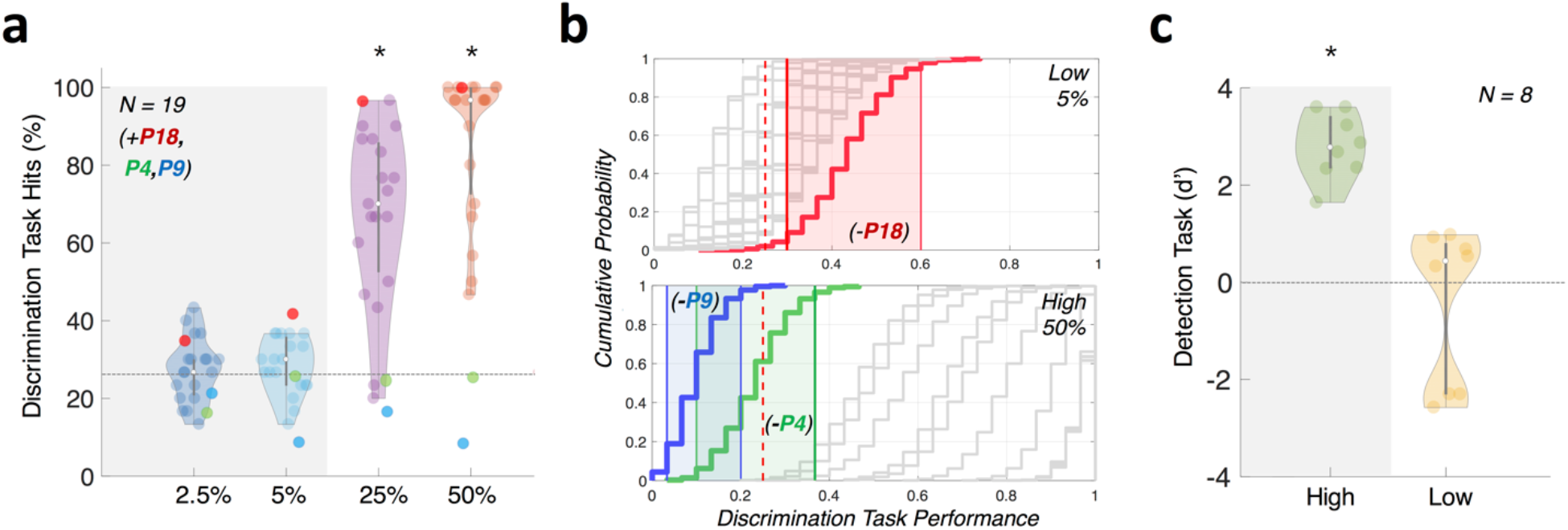
Results from follow-up psychophysics tasks 1 and 2 confirming motion (in)visibility. **(a)** Direction discriminability follow-up psychophysics task 1 for participant exclusion. Accuracy in this task was used to confirm that the overall direction of motion at 5% coherence was not discernible, whilst 50% coherence motion was perceived and reported. Grey and coloured dots represent included and excluded participants, respectively, for the further EEG analysis, based on our bootstrapping analysis (panel b). **(b)** We confirmed our participant exclusion using bootstrapping of performance (plotted as cumulative histogram; included participants plotted in grey and excluded participant plotted in colour). Participants were excluded if they performed above chance for 5% coherence (left panel, N = 1; P18, red) or below chance for 50% coherence (right panel, N = 2; P4, green and P9, blue). Coloured shading indicates 95% confidence interval for the respective excluded participants. Red dotted line indicates chance-level performance. Removed participants are included in panel (a) only as a reference. **(c)** Motion change detectability (d’) for individual participants in psychophysics task 2. We found low-coherence (5%) d’ was not significantly greater than zero but d’ for high-coherence (50%) was. Note: Error bars show standard deviation.

Based on our bootstrapping analyses, we confirmed the visibility and invisibility of the high- and low-coherence motion direction changes, and excluded two participants due to criterion 1 and one due to criterion 2. Due to technical error (corrupted data files), we could not perform bootstrapping analysis for four more participants and we conservatively chose to remove the EEG data of these four participants from further analysis as well.

In summary, we used the data from N=19 participants for ERP, source, and DCM analysis.

#### Follow-up psychophysics task 2: Detectability of motion direction changes

Finally, as another follow-up experiment, we ran a detection task to exclude the possibility that participants were able to ‘sense’ the change of motion direction even in the absence of the perception of motion direction per se, but based on the metacognitive ‘feeling of change’ (Rensink, 2004). To test this, we repeated one block from the main experiment; after removing the letters for the 1-back task but leaving the small, black central circle (Figure 1). The direction of motion and coherence levels were the same as the main experiment. We instructed participants to press the spacebar as quickly as possible whenever they detected a change in the global motion direction of the cloud of dots. Participants were instructed that even if they were unsure, they should report a feeling of change. We regarded a change of global motion direction within a coherence level (which was either 5% or 50%) as a target event. There were 20 events per coherence level. To analyze this behavioral data, we computed the sensitivity measure of d’ using the same signal detection theory method as our main experiment.

### Behavioral results psychophysics task 2: *Confirming lack of ‘feeling’ of change*

We found that d’ was significantly above zero for high-coherence motion direction changes (Figure 2c; one-sided one sample t-test, *M* = 2.77, *SD* = 1.68, *p* > 0.001, *df* = 7) but not for low-coherence motion direction changes (one-sided one sample t-test, *M* = −0.47, *SD* = 1.60, *p* = 0.781, *df* = 7). We took this as evidence that this subset of participants could not even sense direction changes at low-coherence, while they were able to do so at high-coherence.

### EEG preprocessing

We used SPM12 (http://www.fil.ion.ucl.ac.uk/spm/) to pre-process the data. We first re-referenced the raw EEG recordings to the average of all electrodes, down-sampled to 200 Hz and high-pass filtered above 0.5 Hz. We epoched within a trial time window of 100 ms before to 400 ms after the onset of each motion trial. We then removed eyeblink artefacts using the VEOG channels with a bad channel maximum rejection threshold of 20% and by thresholding all channels at 100uV. In the following EEG analysis, we present data without the removal of trials in which a target for the 1-back task occurred or a button response was made. Given that the task and button presses were uncorrelated with the conditions of interest (i.e. motion trials), these are unlikely to have an effect on our results (which we confirmed; data not shown). To obtain the mean ERP per participant per condition, we averaged the epoched data using robust averaging (Wager et al., 2005) with a built-in function in SPM12. This robust averaging process down-weighted each time-point within a trial according to how different it was from the median across trials. Next, because high frequency noise can be introduced during the robust averaging process, we further low-pass filtered the processed data at 40 Hz. Finally, we baseline corrected the data for each participant and condition by subtracting the (robustly averaged) ERP between −100 to 0 ms.

### ERP analysis

We analysed ERPs corresponding to each of the experimental conditions across participants (as well as the visual PE, in the form of the ‘vMMN’: derived from subtracting the standard waveform from the deviant waveform within a condition). We combined channel clusters over left anterior (FC3, F1, F3, F5, AF3), right anterior (FC4, F2, F4, F6, AF4), left central (C3, C5, CP1, CP3, CP5), right central (C4, C6, CP2, CP4, CP6), left posterior (P7, PO3, PO7, PO9, O1) and right posterior (P8, PO4, PO8, PO10, O2) electrodes (see Jack et al., 2015). The significant vMMN time periods were those where the difference waves mean amplitude was below zero across participants (after FDR correction for multiple comparisons at each time point).

### Spatio-temporal statistical maps

We obtained spatiotemporal images of the ERP for each participant and condition across the scalp within the window of −100 to 400 ms. To obtain one 2D image, for each of the 101 time bins, we projected the EEG electrode locations onto a plane and interpolated the electrode locations linearly onto the 32 x 32 pixel grid. We then stacked each 2D image over time to obtain a 3D volume (32 x 32 x 101) and smoothed with a 3D Gaussian kernel of full-width at half maximum: 12 mm x12 mm x 20 ms.

The 3D spatio-temporal image volumes were analysed, on a participant-by-participant basis, with a mass univariate general linear model (GLM) as implemented in SPM12. We estimated the main effects of surprise (i.e., standards vs deviants) and coherence (i.e., high-vs low-coherence), their interaction, and contrasts between standards vs deviants separately within the high- or low-coherence conditions using between-subject F-contrasts. All spatio-temporal effects are reported at a threshold of *P*<0.05 at the cluster-level with family-wise error (FWE) correction for multiple comparisons over the whole spatio-temporal volume. F-statistics are reported as the maximum at the peak-level.

### Source reconstruction

We obtained source estimates on the cortical mesh by reconstructing scalp activity with a single-sphere boundary element method head model, and inverting a forward model with multiple sparse priors (MSP) assumptions for the variance components under group constraints (Friston et al., 2008). One benefit of using MSP for the source reconstruction process is that through the use of empirical Bayes one can select multiple sources to build covariance components for both sparse or distributed source configurations (Friston et al., 2008). Because our main effects of coherence and surprise (and their interaction) spanned the entire epoch we decided to consider the whole peristimulus time window (0 to 400 ms) in the MSP procedure. This allowed for inferences on the most likely cortical regions that generated the sensor-level data across the entire trial time window. We obtained volumes from these reconstructions for each of the four conditions for every participant. These images were smoothed at full-width at half maximum: 12×12×12 mm^3^. We then computed the main effects of (1) coherence, (2) surprise, (3) the interaction (coherence x surprise), as well as the (4) high- and (5) low-coherence PE (standards vs. deviants) using conventional SPM analysis between-subject F-contrasts.

### Dynamic Causal Modelling

We used DCM to estimate a generative causal model that most parsimoniously explained the observed ERPs at the selected source locations with minimal model complexity (Friston, 2003). It is important to note, here, that DCM is a statistical inference of causal models, and ‘causality’ is not established through perturbation or other intervention (Pearl, 2000). We used a data-driven approach combined with a priori locations drawn from the visual motion processing literature, to explain best the observed PE related signals for visible and invisible motion changes. The data-driven spatial location of left inferior temporal gyrus (ITG) was selected based on our source-level analysis. We also included the sensory input nodes of bilateral primary visual cortices (V1) and middle temporal cortex (MT+), which are likely to be the initial cortical stages for visual motion processing (Born & Bradley, 2005), and the bilateral posterior parietal cortices (PPC), which are known for higher-level visual motion processing (Anderson, 1989; Ilg et al., 2004). In total, we assumed seven sources: bilateral V1, MT+, PPC and left ITG (see Results for Montreal Neurological Institute (MNI) coordinates).

We connected our candidate nodes using the same architecture and then exhaustively tested all possible combinations for the direction of modulation(s) amongst these nodes (with one exception, see below). Our model architecture comprised: (1) recurrent (forwards and backwards) connections from V1 to MT+ and MT+ to PPC within the left and right hemispheres, and left MT+ to left ITG, (2) each node was laterally connected with the corresponding node in the other hemisphere (e.g. left V1 laterally connected to right V1), (3) intrinsic (or within-region) modulation only at V1 and MT+, which are known to strongly adapt to repeated visual motion in the same direction (Kohn & Movshon, 2004), and (4) left ITG and left PPC laterally connected. We decided to assign the equal hierarchical relationship between ITG and PPC due to the inconsistent relationship between them reported in the literature (e.g., DeYoe & Van Essen, 1988; Felleman & Van Essen, 1991).

Next, although there could be a huge number of possible DCM modulations based on our 7 identified nodes, we decided to reduce the possible space for modulations based on anatomical information as much as possible. By anatomical criteria, we decided to examine models that always contained recurrent (forwards and backwards) modulation between the input sources of bilateral V1 and MT+ and intrinsic modulations at these input nodes; we refer to this as the ‘minimal’ model (i.e. a base model for all other models). Using this reduced number of potential modulation directions, we were left with 7 connections that could be modulated beyond the minimal model: 4 connections between MT+ and PPC (forwards and backwards in each hemisphere), 2 connections between MT+ and IT (forwards and backwards in left hemisphere) and 1 connection between PPC and IT (one lateral connection in left hemisphere). Note, that we never modulated the lateral connections between the hemispheres. Thus, in total, we tested 129 models (i.e. 2^7^ = 128 + a null model with no modulation at or between any nodes) comprising all combinations of modulation directions (see Figure 7a).

We performed DCM analyses to estimate the parameters of each model, separately for each effect of interest: (1) the visible PE and (2) the invisible PE. For the visible PE, we used the between-trial effect (condition weights) of [0, 1] for the high-coherence standard and deviant trials, respectively. For the invisible PE, we used the weights of [0, 1] for the low-coherence standard and deviant trials, respectively. DCM analyses started from estimation of the log model evidence for each of our 129 generative models fitted for every participant. The log model evidence quantified how well each model could generate ERPs similar to the observed (pre-processed) ERPs for that participant. Importantly, the iterative free-energy minimization process used to calculate the log model evidence penalised models with greater complexity to avoid overfitting (Friston et al., 2008). Once estimated, we used a random-effects (RFX) approach to determine the winning model across participants via Bayesian Model Selection. RFX assumes that a (potentially) different model underpins each participant’s responses; making it robust to outliers and best suited for studying perceptual processes whose underlying network structure is likely to be varied across participants (Stephan et al., 2009). To compute a weighted average of the parameter estimates across all models, we employed Bayesian Model Averaging (BMA) (Penny et al., 2010). BMA weights the estimated parameter with the probability of each model associated with that parameter for all participants. In this way, all models contributed to the final connectivity estimate, with the most probable model having the greatest weight and the least probable model contributing the least to the final estimates (Stephan et al., 2010). In a follow-up DCM analyses, we compared the visible and invisible PE by modelling their interaction using the between-trial effect (condition weights) of [0,1] to contrast the mean ERPs for the visible PE and invisible PE per participant.

## RESULTS

### No difference in distraction by visible or invisible motion directions

After exclusion based on the follow-up psychophysics task, we analysed the performance of the remaining N = 19 participants in the 1-back task to check if they appropriately focused on the central letter task and ignored the background motion during the EEG session of the main task. Based on prior studies (e.g., Tsushima et al., 2006), we expected that our low-coherence background motion would distract participants more than high-coherence motion to degrade the performance of the 1-back task.

Between the high- and low-coherence conditions, we observed a difference in the percentage of hits and false alarms but no difference in d’ or reaction times. The proportion of hits during the high-coherence condition (M = 69.54%, SD = 11.68%) was significantly lower than in the low-coherence condition (M = 73.87%, SD = 12.34%; two-tailed paired t-test, *t*(18) = −3.27, *p* = 0.004; Figure 3a). Mean number of false alarms across participants was also significantly lower in the high-coherence condition (M = 4.95, SD = 4.71) than the low-coherence condition (M = 6.95, SD = 5.34, two-tailed paired t-test, *t*(18) = −2.39, *p* = 0.028; Figure 3b). Combining the proportion of hits and false alarms, we computed the d’ measure. Using this measure, we found no significant differences between d’ for performance on the 1-back task during the high- (M = 3.32, SD = 0.49) and low-coherence conditions (M = 3.31, SD = 0.49, two-tailed paired t-test, *t*(18) = 0.09, p = 0.931) (Figure 3c). Furthermore, we found no differences in the reaction times between the high (M = 557 ms, SD = 59 ms) and low (M = 555 ms, SD = 64 ms, two-tailed paired t-test, *t*(18) = 0.722, *p* = 0.48) coherence conditions (Figure 3d).

**Figure 3.**
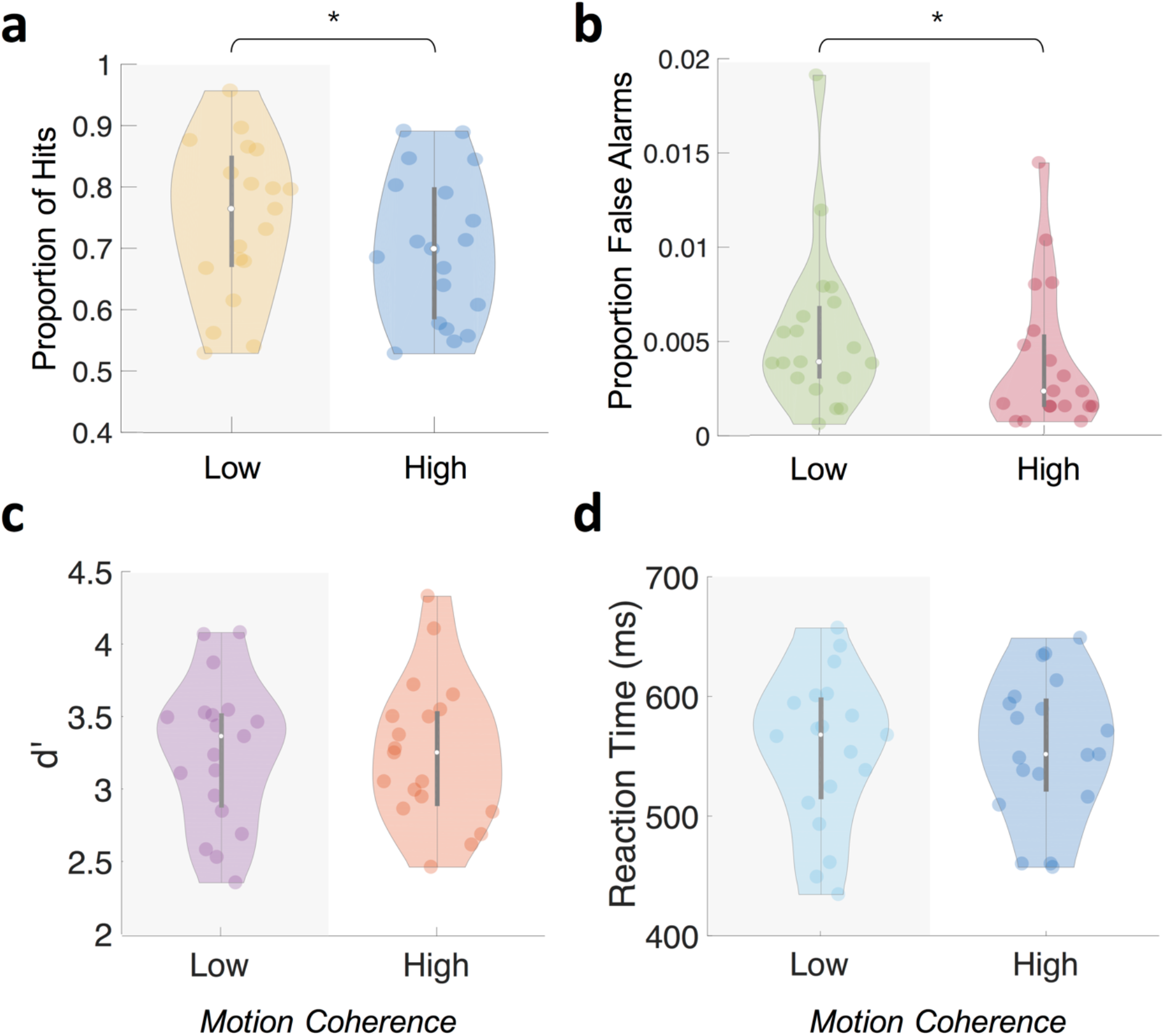
Behavioural results from central 1-back task during Main Experiment. **(a, b)** Mean proportion of hits (a) and false alarms (b) was significantly higher (*p* < 0.01 and *p* < 0.05, respectively) when background motion was low-coherence than high-coherence. (c, d) Mean d’ (c) and reaction times (d) showed no significant difference (*p* > 0.05) between the coherence levels.

Overall, we did not find the results to be consistent with our expectation of greater distraction effects during low-coherence motion trials. We suggest this may be due to the differences between our study and that by Tsushima et al. (2006; such as the different behavioural task and the inclusion of the PE generating stimulus sequence), leading to lower task difficulty and comparable distractibility effects between the two coherence levels. We interpret these results as: (1) participants were able to maintain their focal attention to the 1-back task, (2) participants did not trade off speed for accuracy differently between the two coherence conditions and (3) that the behavioral effects of high- or low-coherence motion were comparable.

### Scalp-level ERP analysis

#### Non-conscious prediction errors occur earlier than conscious prediction errors

We extracted the ERPs from scalp-level electrode clusters to examine the vMMN for visible and invisible PE responses (defined as sustained negativity of vMMN difference waves after correcting for multiple comparisons at each time point, *q* < 0.05). We found that vMMN was evoked from both visible and invisible motion direction deviants. For visible PE responses (Figure 4a and 4b), we observed vMMN (p_FDR_ < 0.05) at left central and anterior channels clusters between 285-295 and 275-400 ms, respectively. For the invisible PE responses (Figure 4c), we observed vMMN at left posterior channels at 150 ms.

**Figure 4.**
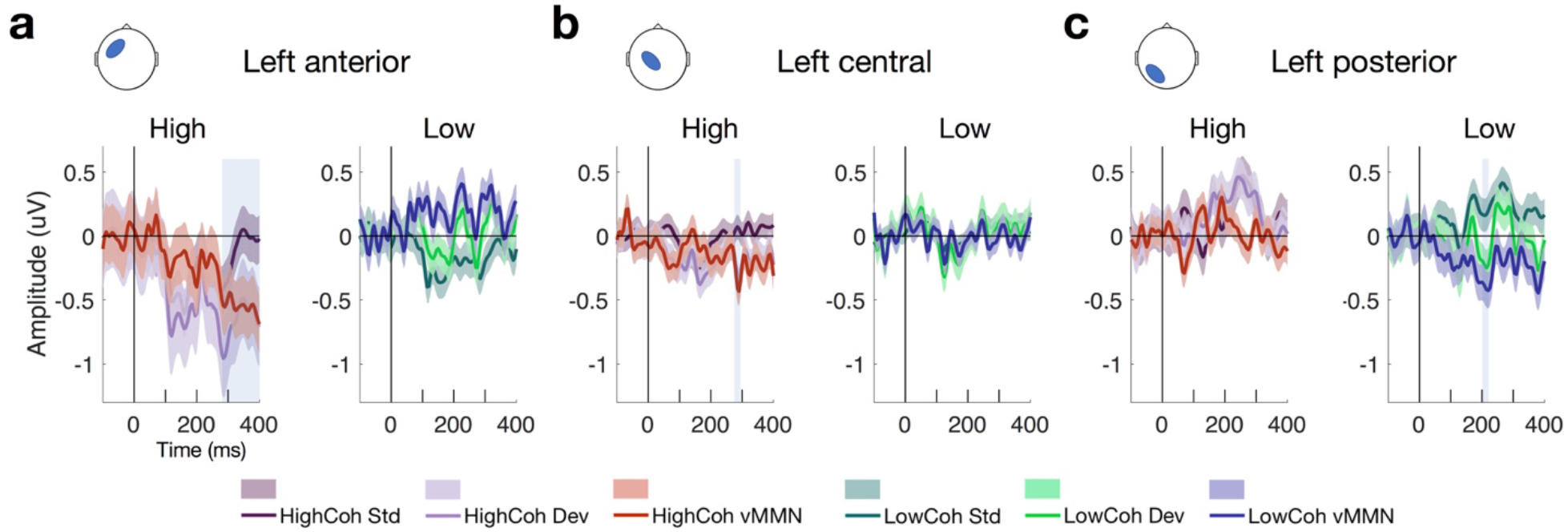
ERP waveforms showing vMMN. Mean ERPs from electrode clusters with significant effects at (a) left anterior, (b) left central and (c) left posterior clusters, showing standard error across participants (N = 19). Left-hand panels show high-coherence ERPs for standard (dark purple), deviant (light purple) and vMMN difference waves (red; deviants - standards). Righthand panels show low-coherence ERPs for standard (dark green), deviant (light green) and vMMN difference waves (blue; deviants - standards). Significance after FDR correction for multiple comparisons is denoted by the shaded blue boxes.

In addition to examining ERPs at the scalp, we used spatio-temporal statistical map analysis in SPM (see Methods). Here, we obtained the 3D images interpolated from ERP data recorded at the scalp and applied between-subject F-tests to quantify the effect of surprise (i.e., prediction error (PE)) within the high- or low-coherence conditions (i.e. visible and invisible PE). Importantly, we found that PEs were evoked from both visible and invisible motion direction deviants. PE to visible motion direction changes (Figure 5a) disclosed a number of significant clusters, ranging from 290 – 395 ms observed across widespread channels. The earliest cluster at 290 ms was found at the central channels (peak-level F_max_ = 25.44, cluster-level p_FWE_ = 0.024) followed by a cluster at left front-temporal and central channels at 380 ms (peak-level F_max_ = 34.12, cluster-level p_FWE_ < 0.001) and at 395 ms (peak-level F_max_ = 67.29, cluster-level p_FWE_ < 0.001). Compared to visible PE, invisible PE occurred earlier and were less spatially spread (Figure 4b); only at 160 ms in left parietal channels (peak-level F_max_ = 25.65, cluster-level p_FWE_ = 0.024).

**Figure 5.**
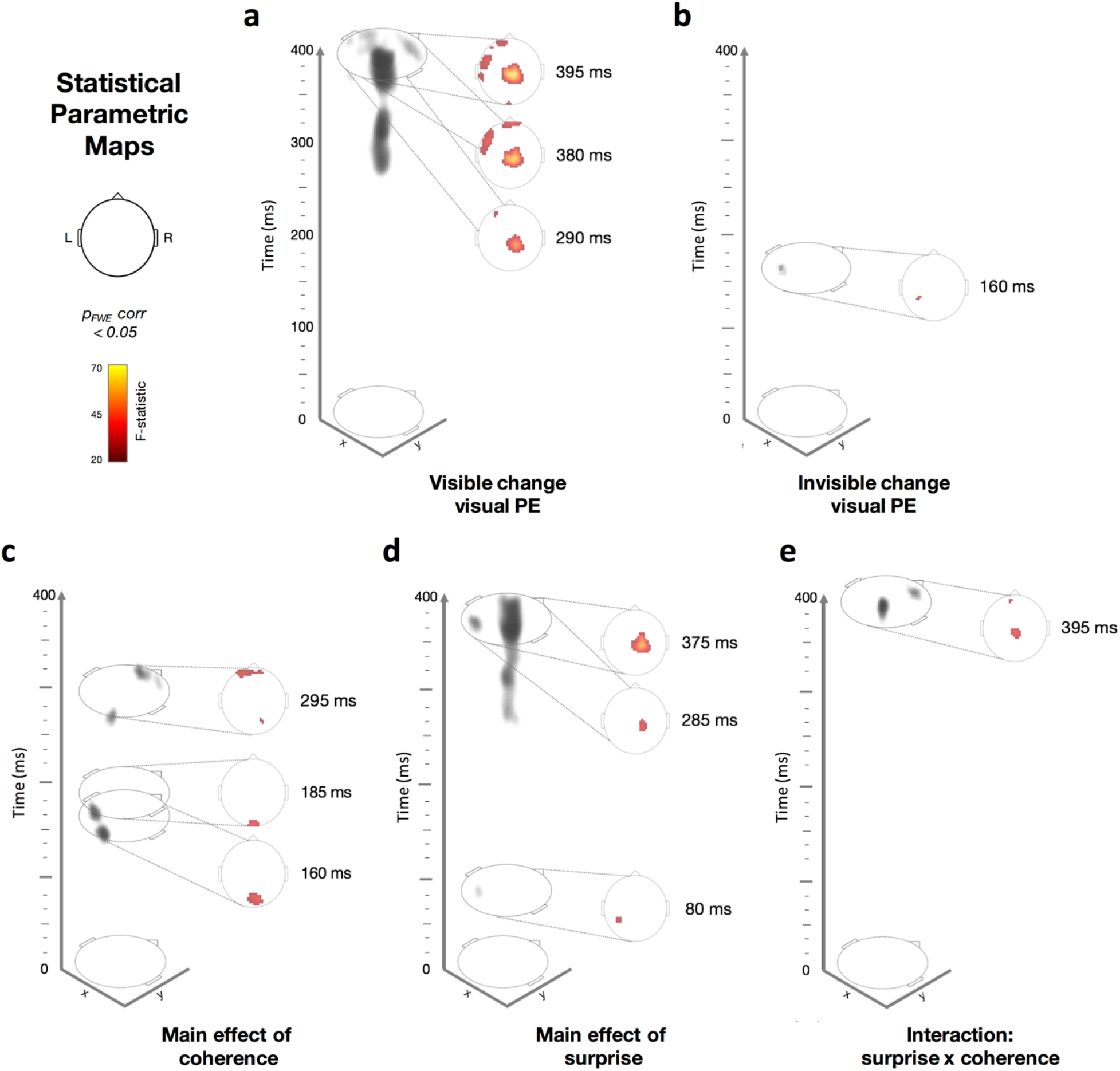
Spatiotemporal Statistical Parametric Maps showing thresholded F-statistic contrasts from the ERP data from (N=19; p_FWE_ corrected < 0.05). Scalp-topography orientation with respect to x- and y-axis is shown as a reference for panels (a-e) The colourmap indicates the range of F-statistic values used in panels (a-e), ranging from 20 (black) to 70 (yellow), where F=24.18 corresponds to p<0.05 with family-wise error (FWE) correction. On the left side of each panel, we show the 3D representation (grey shading) of the significant clusters across space (x, y) and time (z, from 0 to 400ms after the trial onset). On the right side within each panel, we show a time slice (e.g. Z=395ms, for example) of the F-statistics. Dotted lines connect the time slice with its location within the 3D volume. (a) Visible standards versus (vs) deviants. (b) Invisible standards versus (vs) deviants. (c) The main effect of coherence (high vs low; regardless of surprise) (d) The main effect of surprise (standards vs deviants; regardless of coherence). (e) An interaction between coherence and surprise. (p<0.05).

Next, we applied between-subject F-tests to quantify the main effects of coherence (Figure 5c), surprise (Figure 5d) and their interaction (Figure 5e). The main effect of coherence (Figure 5c) disclosed three significant clusters. The first cluster at 160 ms was located occipitally (peak-level F_max_ = 34.37, cluster-level p_FWE_ = 0.002), the second at 185 ms occurred in the same location (peak-level F_max_ =35.72, cluster-level p_FWE_ = 0.004) and the third at 295 ms was found at right occipito-parietal and frontal channels (peak-level F_max_ = 32.31, cluster-level p_FWE_ = 0.001). The main effect of surprise (Figure 5d) showed three significant clusters. The first at 80 ms occurred at left parietal channels (peak-level F_max_ = 29.53, cluster-level p_FWE_ = 0.014), the second at 285 ms was located at right central channels (peak-level F_max_ =34.66, cluster-level p_FWE_ < 0.001) and the third at 375 ms was observed in the same location (peak-level F_max_ = 55.29, cluster-level p_FWE_ < 0.001). Finally, we observed an interaction between surprise and coherence (Figure 5e) at central and frontal channels at 395 ms (peak-level F_max_ = 35.93, cluster-level p_FWE_ = 0.002).

### Source-level analysis

#### Left ITG as a source for conscious PE

We applied MSP source reconstruction to estimate the cortical regions involved in generating PE to visible and invisible motion direction changes. In Figure 5 (the scalp-level maps), we found that the main effects of surprise, coherence, and the interaction spanned the whole epoch, thus we decided to use the whole epoch data (0-400 ms) rather than to temporally constrain the data for source reconstruction (see Methods for details). Figure 6 shows the significant source-level results for the main effect of surprise and PE to visible changes at an uncorrected threshold of *p*<0.001. We did not find any significant sources for invisible PE, main effects of coherence or interaction when we corrected for multiple comparisons. For the main effect of surprise, we found one significant cluster in the left ITG ([−52 −26 −30, peak-level F_max_ = 22.19, cluster-level p_FWE_ = 0.003, Figure 6a). For the PE to visible changes, we found a similar cluster in the left ITG ([−48, −12, −32], peak-level F_max_ = 17.91, cluster-level p_FWE_ = 0.039, Figure 6b). The effect of surprise for the invisible conditions revealed a cluster on the right hemisphere (Figure 6a) which did not survive correction.

**Figure 6.**
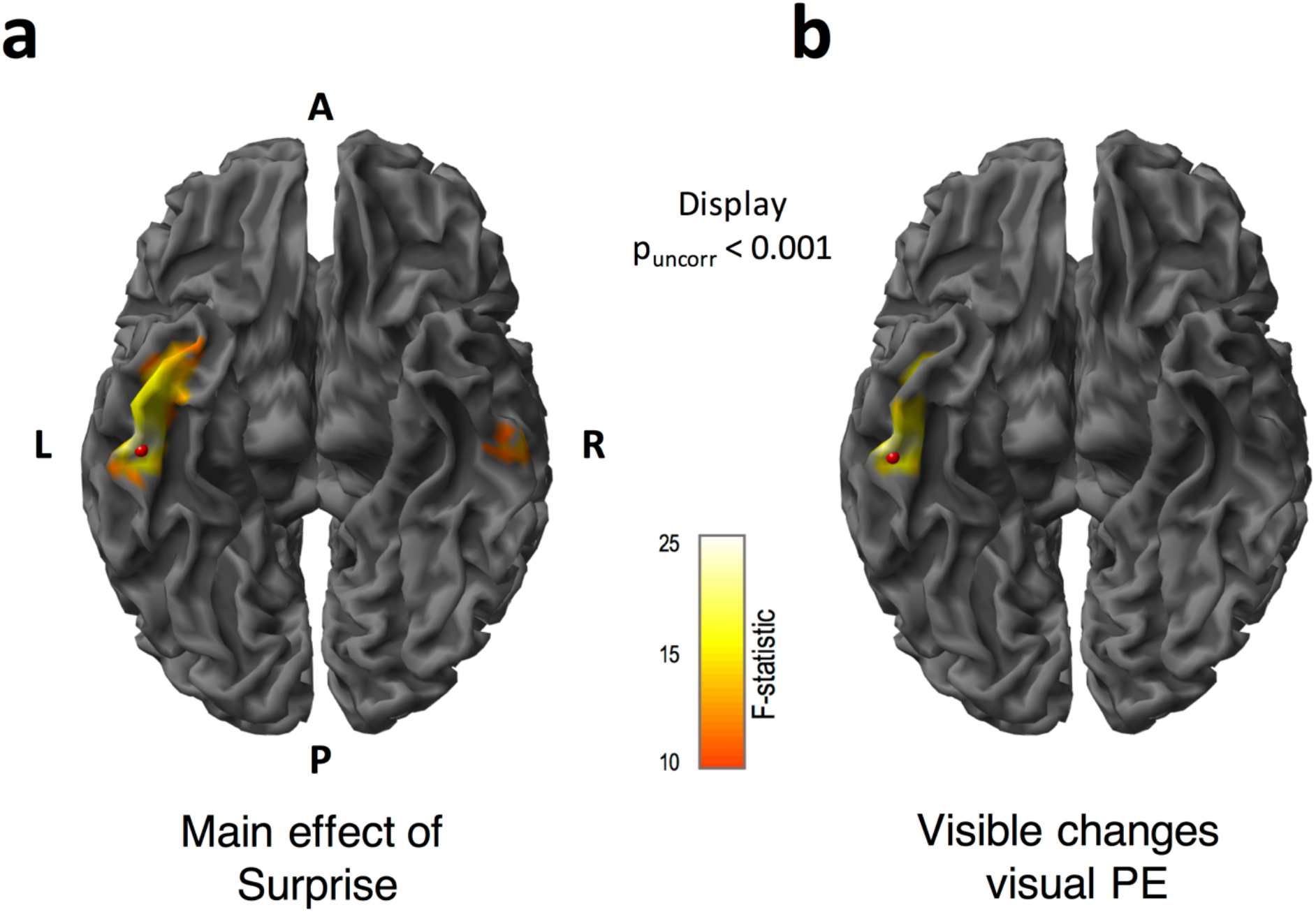
Source reconstructed F-statistics for the main effect of surprise, and prediction errors (PE) to visible motion direction changes. We obtained source locations using multiple sparse priors (MSP) source reconstruction. These maps are displayed at p_uncorr_<0.001. (a) For the main effect of surprise, we found a significant cluster in the left inferior temporal gyrus. (The cluster in the right inferior temporal gyrus did not survive after correction). (b) Source localisation for visible motion changes revealed a significant cluster in the left inferior temporal gyrus. The F-statistic is displayed for all contrasts within the range of 10 to 25, where F=12.11 corresponds to p<0.05 FWE corrected. A = anterior, P = posterior, L = left and R = right. Red beads indicate the location of the peak of F-statistic

### Dynamic Causal Modelling

We used DCM to examine how the source location identified in the previous step, interacted with cortical locations known to specialize in motion processing, and how the strengths of these interactions are modulated by the visibility of motion changes. Specifically, we identified one region at the FWE corrected (*p*<0.05) threshold (between-subject F-tests) (Anatomy Toolbox; Eickhoff et al., 2005): the left **ITG** (MNI coordinate: [−48, −12, −32], Figure 6). We included the sensory input nodes of bilateral **V1** (MNI coordinates: left [−14, −100, 7] and right [17, −97, 9]) and bilateral **MT+/V5** (MNI coordinates: left [−48, −69, 7] and right [50, −66, 11]) as these regions are essential for visual motion processing (Born & Bradley, 2005; Plomp et al., 2015). We also included the bilateral **PPC** (MNI coordinates: left [−46, −46, 54] and right [52, −42, 50]) because these regions are known for higher-level visual motion processing (Ilg et al., 2004). Based on our 7 identified nodes, we tested 129 models comprising all combinations of modulation directions (building on the fixed ‘minimal model’; see Methods for more information). Figure 7a shows the defined model space, including: the fixed model architecture (white arrows) and how each model was allowed to vary in terms of the direction of modulation (black arrows).

**Figure 7.**
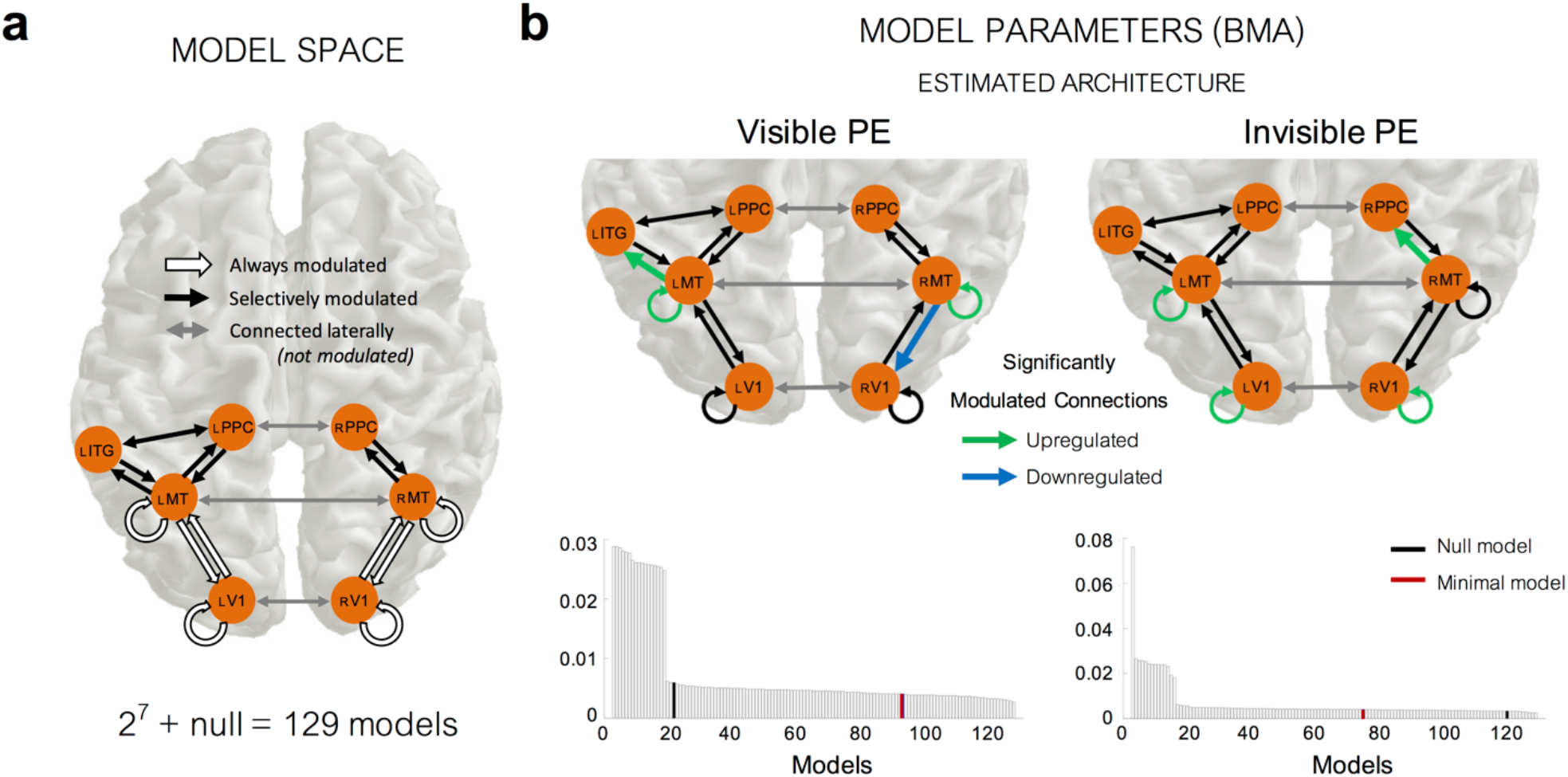
Dynamic Causal Modelling (DCM) for visible and invisible PE responses. (a) Model space showing the architecture used in all models and modulation of these connections. We used the same architecture in all models, what differed were the allowed modulations of the connection strength between nodes. We systematically varied the modulation direction of all black connections in a binary manner to obtain 2^7^ = 128 models (+ a “null model” which did not have any modulation at or between any nodes). One of the 2^7^ = 128 models also contained “the minimal model”, in which the only modulated connections were those between V1 and MT, and intrinsic modulation within V1 and MT. (b) Model evidence results from our random-effects (RFX) Bayesian Model Selection analyses, displayed as the exceedance probability of each model (i.e. the probability that a particular model is more likely than any other model given the group data). Using Bayesian Model Averaging (BMA) we were able to obtain weighted parameter estimates using the model evidence across all models and participants. Significantly modulated connections (False Discovery Rate corrected) are shown in green (upregulated) and blue (downregulated).

### Optimized DCM modulation parameters explaining differential ERPs for visible and invisible PE

Once the log model evidence for every model and participant was estimated, we used RFX Bayesian Model Selection to determine the winning model at the group-level. Figure 7b shows, for both the visible and invisible PE, the exceedance probability of all models at the group-level. The exceedance probability is the probability that a particular model is more likely than any other model given the group data.

For the visible PE, there was no single clear winning model amongst our 129 models. This is unsurprising when using RFX analysis as effects can be diluted if a large number of models are compared in a small sample size. In this situation, it is recommended not to single out the best architecture alone, but rather to use BMA to consider all models, taking into account the strength of the evidence for each model. After applying BMA, we found significant positive modulation of the intrinsic self-connections of left MT+ (mean parameter outputs = +0.2004, two-tailed t-test against 0: *pFDR*= 0.0486, *df* = 18), and right MT+ (+0.2644, *pFDR* = 0.0011, *df* = 18). This result confirms what we expected from the nature of our oddball paradigm; the visible change induced PE relies, in part, on intrinsic (self) modulation at MT+ due to a release from adaptation (i.e. repetition suppression) following the onset of deviant motion. In addition to this, we also found two more significantly modulated connections. We found a *backward* connection from right MT+ to right V1 was significantly downregulated (−0.1310, *pFDR* = 0.0453, *df* = 18) and a *forward* connection from left MT+ to left ITG was significantly positively modulated (+0.0875, *pFDR* = 0.0060, *df* = 18).

For the invisible PE, as with our visible PE DCM analyses, in the absence of one single winning model, we again applied BMA finding that, similarly to the visible PE DCM results we observed significant positive modulations for intrinsic modulations within left V1 (+0.1501, *pFDR* < 0.0001, *df* = 18), right V1 (+0.1400, *pFDR* = 0.0081, *df*= 18) and left MT+ (+0.2821, *p FDR* < 0.0001, *df*= 18), reflecting a release from adaptation upon a change in motion direction, despite no awareness of this change occurring. One difference, compared to the visible PE, was the direction of one other significantly positively modulated connection *forwards* from right MT+ to PPC (+0.2048, *pFDR* = 0.0094, *df* = 18). We found no significantly modulated backward connections within our DCM analyses of the invisible PE.

Finally, we directly compared differences in effective connectivity between the visible and invisible PE by modelling their interaction. We applied RFX BMA to determine the model parameters in the absence of one single winning model and found two significant connections (prior to FDR correction) between left ITG and left PPC and at a self-connection of left MT+. However, these connections did not survive correction for multiple comparisons. We interpret these findings as weak but consistent with our source results showing left-hemisphere localisation of the visible PE.

## DISCUSSION

In this study, we aimed to elicit PE related neural activity via changes in stimulus statistics that were consciously visible or not. As planned, we successfully manipulated the awareness of the stimulus changes using supra- and sub-threshold motion coherence of a dynamic random-dot display and confirmed the visibility of the motion direction changes with a follow-up psychophysics task. Our ERP analyses (Figure 4 and 5) confirmed robust visible PE responses, which were widespread in space and time and invisible PE responses, which were more confined in space and time. Our source level analysis located the source of the visible PE response in the left ITG. Using the identified location of the left ITG and the cortical nodes known to be critically involved in motion processing (V1, MT+, and PPC), we performed DCM analyses, hoping to reveal the network underpinning visible and invisible PE responses.

### Confirmation of motion direction (in)visibility

It is important that we first acknowledge that our method of rendering our stimuli invisible (or weaker stimulus paradigms in general, e.g. Dehaene, 1999; Dehaene & Naccache, 2001; Gaillard et al., 2009, Del Cul et al., 2007, van Vugt et al., 2018) confounds the issue of conscious perception with the issue of stimulus change. We also point out that alternative approaches which eschew this issue (such as binocular-rivalry like paradigms), do suffer from other issues such as the effects of the report. That is, when the physical stimulus input is equated, the effects of reports can be falsely interpreted as the neural correlates of consciousness (Frässle et al., 2014, Tsuchiya et al., 2015, Wilke et al., 2009). Thus, we foresee the combination of both approaches (Tsuchiya et al., 2016) will be important for future studies.

Secondly, we must highlight the factors leading to the design of our follow-up psychophysics task 1 that was used to confirm (in)visibility of motion direction at high- and low-coherence. The design of this task was motivated by a number of concerns with presenting motion stimuli identical to that used in the main experiment. We opted to change the motion parameters during this task primarily because of our concern on perceptual learning and habituation during the visibility check. To alleviate this issue, we made four critical changes in the stimulus design: we lengthened the visual motion presentation time, removed the distracting central letter task, intermixed 4 coherence levels, and reduced the direction alternatives. All of these changes, in addition to the fact that during the main experiment participants did not pay attention to the motion, should have made our visibility assessment rather conservative; that is, we should have detected any participants who were aware of the low-coherence visual motion direction changes during the main task using this visibility task. Furthermore, while our choice of 30 trials was on the smaller end in terms of the number trials as a visibility check, we performed non-parametric bootstrapping analysis on an individual basis, which is the most sensitive method to detect aware participants. Overall, we are confident in our method of confirming that the motion direction changes were visible or invisible.

### Temporal features of the prediction error (PE) responses with visible and invisible motion changes

We were able to elicit PE responses for both visible and invisible motion direction changes. Scalp ERPs (Figure 4) disclosed a significant early component at 150 ms for invisible PE (i.e. the low-coherence vMMN difference wave) at left posterior channels. In contrast, we observed two significant later components for the visible PE (high-coherence vMMN difference waves) between 285-295 and 275-400 ms at left central and left anterior clusters, respectively. Our spatio-temporal scalp-level statistical mapping analysis (Figure 5) supported these findings. Here, after FWE correction, PE evoked by invisible motion direction changes peaked at parietal channels earlier in time (<160 ms) than PE to visible motion direction changes (>290 ms), which were observed at central and fronto-temporal channels.

Our finding is consistent with some studies that elicited non-conscious visual PE. In an MEG study using vertical gratings and backward masking of rapidly presented deviants, Kogai et al. (2011) found similar latencies in the non-conscious PE response from 143-154 ms in striate regions. Further, Czigler et al., (2007), backward masked coloured checkerboards at different stimulus onset asynchronies (SOAs) and found mask SOAs at 40 and 53 ms elicited PE between 124-132 ms at occipital channels (but that participants had some level of awareness of the deviant stimuli). Other studies, however, report later PE to invisible oddballs. For example, Jack and colleagues (2017) found slightly later non-conscious PE (to deviant stimuli presented monocularly to the non-dominant eye) that peaked around 250 ms (as did the conscious PE). This longer latency for the non-conscious PE was also echoed in another study from the same group (Jack et al., 2015). We suggest the discrepancies in the latencies arise from both stimulus differences and in the methods used to render stimuli non-conscious. Future paradigms in which both the standard and deviant stimuli can be rendered invisible may help disentangle conscious and non-conscious PE effects.

But why does the invisible PE occur earlier than visible PE? Here we offer two possible explanations. The first is related to the effects of attention. Previous visual PE studies that have used consciously *perceivable motion direction* changes found an earlier PE at 142-198 ms regardless of the attentional load on the irrelevant task (Krëmlacek et al., 2013). A separate study found a later PE component emerged when attention was directed to the PE generating stimuli (Kuldkepp et al., 2013). Indeed, in a binocular rivalry vMMN study (using grating stimuli), van Rhijn et al. (2013) observed two early components from 140 to about 220 ms both when attention was directed to the rivalry stimuli and when attention was diverted. But a second late negativity (270 to 290 ms) was observed only when attention was directed to the rival stimuli. In our paradigm, participants may have attended to the visible motion resulting in a delayed PE. Conversely, invisible motion may not have deployed attention, which resulted in an earlier PE. However, this explanation does not speak to the neural mechanisms. An alternative explanation lies on a putative difference in adaptation (or repetition suppression) for visible and invisible stimuli. It is plausible that stronger adaptation in MT+ to the visible stimuli may mask the earlier PE component in V1. This adaptation effect may be weaker in MT+ for invisible stimuli, although this interpretation does not fully concur with our DCM results showing adaptation in V1 (but not right MT+) for invisible stimuli (see below for further discussion). Additionally, the later PE component for visible stimuli may be more related to the awareness of prediction violation. Further studies that orthogonally manipulate adaptation and prediction might be able to distinguish between these possibilities (Kok et al., 2012; Summerfield & Egner, 2009; Summerfield et al., 2006) and disentangle the two components underlying the (‘classical’) vMMN; namely, the ‘genuine’ vMMN PE response and the effects of neural adaptation (O’Shea, 2015).

### Spatial features of the prediction error (PE) responses with visible motion changes

In terms of spatial characteristics of visible PE, and based on previous studies by Tsushima et al., (2006), we expected to detect prefrontal source activity to our supra-threshold visible motion stimuli. Instead, our source-level analyses (which pooled responses over time) provided evidence that visible PE, as well as the main effect of coherence (regardless of the level of surprise), were generated by cortical sources within the left ITG. We suggest these source-level differences might have arisen from differences, most likely, in our measurement modalities (fMRI in Tsushima et al, 2006, and EEG in our study), as well as differences in experimental paradigms, namely, our inclusion of the roving oddball design instead of coherence differing on a trial-by-trial basis and our 1-back task.

According to our literature search, we note one study of visual consciousness that supports our findings of left hemispheric lateralization (O’Shea et al., 2013). In this study, the authors used binocular rivalry to seek brain activity that could predict visual consciousness and found early activity (around 180 ms) confined to left parietal-occipital-temporal regions. Beyond this, literature on the lateralization of conscious awareness of specific stimuli to the left hemisphere is scarce; reported only in a handful of studies of emotional processing (Gazzaniga, 2000; Kimura et al., 2004; Meneguzzo et al., 2014; Prete et al., 2015; Shepman et al., 2016; Williams et al., 2006). In these studies, subliminally presented face stimuli activated right amygdala (via a subcortical route) in response to fear but left amygdala for supraliminal fear (Williams et al., 2006), and using masked or unattended affective-stimuli activated the right hemisphere in both the visual (Kimura et al., 2004) and auditory domains (Shepman et al., 2016). However, unlike these studies, we used visual motion stimuli and examined the difference between PE rather than responses to visual stimuli more generally. Another possible explanation for the observed lateralization of conscious perception of motion changes to left hemisphere comes from work in split-brain patients showing the left hemisphere is more adept at monitoring probabilities to infer causal relationships based on series of events over time (Gazzaniga 2000; Roser et al., 2005) and at creating internal models to predict future events (Wolford et al., 2000). This suggests that when stimuli are consciously perceivable, the left ITG generates and updates predictions and PE. However, due to the lack of literature, specifically on visual PE lateralization, further work is needed to fully understand the role of left ITG source for conscious PE processing.

### Network level model of causal connections investigated by DCM

Using DCM, we aimed to extend earlier visual PE studies by investigating the network level properties underlying the generation of PE to visible and invisible changes. The primary question of our DCM analysis was whether there was evidence for top-down modulations for PE to visible or invisible change. According to the predictive coding framework, when an unexpected stimulus occurs, the PE signal is propagated from lower to higher brain areas, resulting in upregulation of forward connectivity. This, in turn, is followed by the revised prediction from high-to low-level brain areas, resulting in increased feedback connectivity. Previous studies are consistent with this theory that consciously perceived PE lead to increases in both feedforward and backward connectivity (e.g., Boly et al., 2011). Our finding of increased forward connectivity from left MT+ to ITG for visible PE is consistent with the first part of the theoretical prediction. What is puzzling is that conscious PE was accompanied by significant decreases in top-down feedback connectivity from right MT+ to right V1 (Figure 7b). This means that the neural prediction from MT+ to V1 decreased when the motion direction was unexpected, which appears inconsistent with the general framework of predictive coding. One possible explanation is that when the prediction is violated, the system suspends prediction, corresponding to the down regulated prediction from the high-to low-level area. Invisible PE, on the other hand, only induced enhanced feedforward connectivity from right MT+ to PPC.

Common to both the visible and invisible PE were significant modulations of the self-connections within lower-level visual areas of V1 and MT+. This suggests that both types of PE rely (in part) on a release from adaptation (i.e. repetition suppression) upon deviant motion onset (i.e. change in stimulus statistics). Whether the adaptation effects were observed at V1 or MT+ depended upon whether this change was consciously perceived. That is, visible change PE relied on adaptation at bilateral MT+, whilst invisible change PE relied on adaptation at bilateral V1 (and left MT+). One possible explanation for the stronger effects observed for visible PE at MT+ than V1 could be related to the subcortical visual motion pathway that carries visual information directly from the lateral geniculate nucleus to MT+ (Sincich et al., 2004). It is possible that when the motion signal is more coherent (i.e. stronger), this subcortical pathway is also activated (in addition to that between V1 and MT+), leading to a stronger motion signal in MT+. Subsequently, if MT+ is more highly activated compared to V1, this may lead to greater adaptation effects upon deviant motion presentation. Alternatively, when the motion signal is weak (i.e. low-coherence), this subcortical route is not activated, and thus, V1 and MT+ may have comparable levels of adaptation (as observed in our invisible PE DCMs). Findings that the generators of visual PE to motion changes are located in motion centres or the dorsal pathway itself have been suggested previously by other studies of visible motion direction changes (Kremlacek et al., 2006; Pazo-Lavarez et al., 2004). We add to these findings by showing that this still holds when the changes are invisible.

## CONCLUSION

We provide new insights into the brain mechanisms underpinning visual change detection, even in the absence of awareness, when task reporting is not required. We lend support for visual PE in response to both consciously and non-consciously perceivable changes; with the former evidenced as stronger and more widespread cortical activity. Our findings suggest hemispheric lateralization within the left hemisphere when motion changes were visible. Using DCM, we found that both types of PE were generated via a release from adaptation in sensory areas responsible for visual motion processing. The overall pattern emerging from our study reveals a complex picture of down- and up-regulation of feedforward and feedback connectivity in relation to conscious awareness of changes. To test the generality of our findings, further investigations are necessary, especially with techniques that explicitly manipulate conscious awareness under comparable task conditions testing for the neuronal effects on prediction and surprise.

## Author contributions

ER, NT, & MG designed the study; ER collected and analyzed data; ER wrote the first draft of the manuscript; ER, NT, & MG edited the manuscript.

## Acknowledgements

We thank the volunteers for participating in this study. We also thank both reviewers for their insightful comments, which helped improved the manuscript.

